# Early disruption of neurogenesis and neural architecture by Amyloid-β and Tau during *Drosophila* development

**DOI:** 10.64898/2025.12.31.696993

**Authors:** Khushboo Sharma, Neha Tiwari, Madhu G. Tapadia

## Abstract

Alzheimer’s disease is recognized as a late-onset neurodegenerative disorder; however, accumulating evidence suggests that disease-associated proteins may exert deleterious effects much earlier during neural development. Using a *Drosophila* model expressing human amyloid-β and Tau, we investigated the impact of these proteins on embryonic neurodevelopment and neural stem cell dynamics. Embryos expressing amyloid-β and Tau exhibited pronounced defects in axonal patterning and compromised neuronal cytoskeletal integrity, indicating early disruption of neural architecture. We observed aberrant elevation and mislocalization of the cell fate determinant Prospero, leading to premature neuronal differentiation and disruption of progenitor lineage progression. Consequently, larval brains displayed a reduced number of Prospero-positive ganglion mother cells, reflecting impaired neuroblast lineage maintenance. Cell-cycle analysis using the FUCCI system revealed altered neuroblast cell-cycle dynamics, characterized by depletion of proliferative populations and enrichment of G1-phase cells, indicative of stem cell exhaustion. In addition, increased apoptotic signaling was detected in larval neural tissues expressing amyloid-β and Tau. Together, our findings demonstrate that amyloid-β and Tau disrupt asymmetric division, differentiation, and self-renewal of neural stem cells during development, resulting in reduced neuronal output and defective brain organization. This study highlights that amyloid-β and Tau exert neurotoxic effects early in development, providing mechanistic insight into how neurodevelopmental perturbations may contribute to later neurodegeneration.

## 1. INTRODUCTION

Over 50 million people across globe have dementia and Alzheimer’s disease (AD) contributes to more than 60% of dementia related cases. Alzheimer’s disease (AD) is a progressive neurological disorder with irreversible loss of neuron with aging and associated complications that are major contributor of cognitive impairment and dementia (AD report 2019). AD is regarded as a progressive late-onset neurodegenerative disorder characterized by progressive synaptic loss, neuronal death, and cognitive decline; however, early onset forms are also known (Kamatham et al., 2024; Tahami et al., 2022; Querfurth & LaFerla, 2010). Both the early-onset AD and late-onset AD have included extracellular accumulation of amyloid-β (Aβ) peptides/neuritic plaques and intracellular aggregation of hyperphosphorylated Tau protein/ neurofibrillary tangles, both of which are strongly associated with neuronal dysfunction and neurodegeneration in the adult brain (Gotz et al., 2019; Spillantini & Goedert, 2013 Hardy & Selkoe, 2002;). Also, the early onset has greater Tau burden (Mendez 2019; Palasi et al., 2015). Consequently, most AD research has focused on age-dependent toxicity, synaptic failure, and neuronal loss that occur during adulthood and aging (Zhang et al., 2024).

Emerging evidences however, suggests that Alzheimer’s disease–associated proteins may exert deleterious effects much earlier than previously suggested (Marchetti & Abbracchio, 2005; Mattson, 2004). Increasing clinical and experimental studies indicate that Aβ and Tau can influence neural development, stem cell behavior, and circuit formation, raising the possibility that neurodevelopmental perturbations may predispose the nervous system to later vulnerability (Winner et al., 2011; Haughey et al., 2002). Alterations in neural progenitor proliferation, differentiation, and lineage maintenance have been reported in mammalian models expressing AD-related proteins, supporting the idea that AD pathology may have developmental origins (Mu & Gage, 2011; Winner et al., 2011). Despite these observations, the precise impact of Aβ and Tau on early neurogenesis and nervous system organization remains poorly understood.

In the past years, the *Drosophila* model for neurodegenerative disease research has contributed remarkably into the identification of modifier genes for the pathology. The genetic and functional analyses that have been discovered in *Drosophila* provide insights into research that will reveal pathogenic mechanisms of disease due to conservation of genes and signalling pathways and comparatively rapid genetic analysis (Nitta and Sugie, 2022). Progressive neurodegeneration has been widely induced by human Aβ and Tau in *Drosophila* brain and has been used as a simple *in vivo* model for the molecular pathogenesis of AD (Sharma et al., 2025; Vourkou et al., 2023; Anupama et al., 2022; Jeon et al., 2020). Although, *Drosophila* with no peptides homologous to Aβ, flies expressing human Aβ in all nervous systems have displayed impaired learning, accumulation of amyloid plaques, neurodegeneration, and shortened lifespan, as observed in patients with AD (Crowther et al., 2005; Iijima et al., 2004; Finelli et al., 2004). *Drosophila* embryonic CNS development has also proven to be a useful model for studying the developmental biology due to conservation of signalling pathways (Allan and Thor 2015). Novel insights into issues of human health have also originated from the study of *Drosophila* CNS development (Bolus et al., 2020; Thomas et al., 1988). Many core mechanisms regulating neurogenesis, cytoskeletal organization, and neuronal differentiation are conserved between flies and mammals (Lupo et al., 2021). Importantly, *Drosophila* models expressing human Aβ and Tau have been extensively used to study neurodegeneration, demonstrating synaptic defects, neuronal loss, and behavioural impairments (Crowther et al., 2005; Wittmann et al., 2001). However, most studies have focused on larval or adult stages, leaving the effects of these proteins on embryonic neurodevelopment largely unexplored.

Neurodevelopment is a highly coordinated process involving asymmetric division of neural stem cells, precise regulation of cell-cycle dynamics, lineage-specific differentiation, and establishment of stereotyped neuronal architecture (Homem et al., 2015; Homem & Knoblich, 2012). In *Drosophila*, neuroblasts function as neural stem cells that divide asymmetrically to self-renew while generating ganglion mother cells (GMCs), which subsequently differentiate into neurons or glia (Doe, 2008). This process is tightly regulated by intrinsic factors such as the transcription factor Prospero, which governs the transition from stem cell maintenance to neuronal differentiation (Choksi et al., 2006). Disruption of neuroblast proliferation, cell-cycle progression, or Prospero-mediated fate specification can lead to premature differentiation, stem cell exhaustion, and defective brain development (Homem & Knoblich, 2012). In addition to neurogenesis, proper development of the nervous system requires accurate axonal pathfinding, cytoskeletal integrity, and coordinated interactions between neurons and glial cells. Microtubule-associated proteins, such as Tau and its functional homologs, play crucial roles in maintaining cytoskeletal organization and axonal stability (Tiwari and Tapadia, 2025; Morris et al., 2011; Avila et al., 2004; Hummel et al., 2000). Perturbations in cytoskeletal dynamics during development can result in axonal misrouting, disrupted neural connectivity, and long-lasting defects in brain architecture (van den Heuvel & Pasterkamp, 2008). Whether expression of Aβ and Tau interferes with these processes during early developmental stages remains an important unanswered question.

In this study, we used *Drosophila* models expressing human Aβ and Tau to investigate their impact on embryonic and larval neurodevelopment. By combining lineage-specific genetic manipulation with immunohistochemical analyses, we examined axonal architecture, neuronal cytoskeletal integrity, neural progenitor fate specification, glial organization, neuroblast cell-cycle dynamics, and apoptotic responses. Our findings reveal that Aβ and Tau induce early disruption of neurogenesis and neural architecture, characterized by aberrant axonal patterning, misregulation of Prospero-dependent lineage progression, impaired neuroblast proliferation, and increased apoptotic signaling. Together, this work provides direct evidence that Alzheimer’s disease associated proteins compromise neural development well before the onset of overt neurodegeneration, offering new insight into the developmental contributions to neurodegenerative disease pathology.

## 2. RESULTS

### 2.1 Defective axonal pattering and impaired neuronal cytoskeletal integrity in Aβ and Tau expressing embryos

During embryonic neurogenesis, establishment of a stereotyped axonal scaffold is essential for proper nervous system organization. The microtubule-associated protein, Futsch (recognized by the monoclonal antibody 22C10) is expressed in post-mitotic neurons prior to axonogenesis and marks axons, dendrites, and neuronal cell bodies, making it a reliable indicator of neuronal architecture and cytoskeletal integrity during development (Hummel et al., 2000). We therefore used Futsch immunostaining to assess axonal patterning and neuronal cytoskeletal organization in embryos expressing amyloid-β (Aβ) or Tau.

Expression of Aβ or Tau was driven in either differentiated neurons (*elav-GAL4*) or neural progenitors (*insc-GAL4*) using the GAL4/UAS system. In stage 15–16 control embryos, neuronal architecture was highly organized, characterized by well-fasciculated longitudinal connectives, properly formed anterior and posterior commissures, and a stereotyped axonal scaffold along the ventral nerve cord [Fig. 1A (v–vi); Fig. 1B (v–vi)]. Futsch staining revealed continuous and uniform axonal tracts, indicative of intact cytoskeletal organization. In contrast, embryos expressing Aβ or Tau displayed pronounced defects in neuronal architecture. Futsch immunoreactivity revealed severe axonal disorganization, including defasciculation of longitudinal tracts, discontinuities within axon bundles, and misrouting of both longitudinal and commissural axons [Fig. 1A (xi–xii, xvii–xvii); Fig. 1B (xi–xii, xvii–xviii)]. These defects indicate a disruption of cytoskeletal stability and axonal guidance during embryonic development.

**Figure 1.**
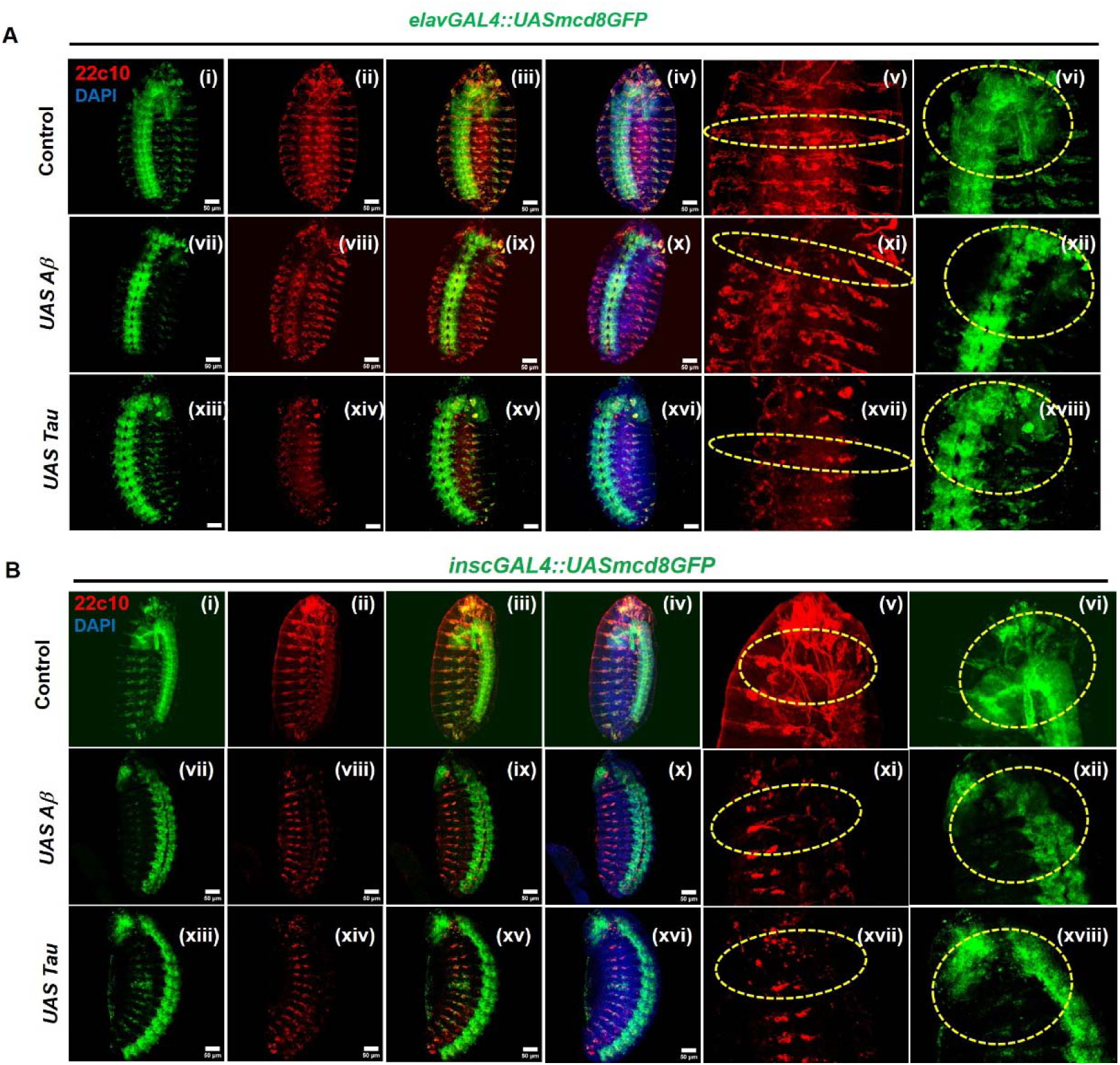
Amyloid-β and Tau disrupt axonal patterning and neuronal cytoskeletal organization during embryonic development. **(A)** Representative confocal images of stage 15–16 *Drosophila* embryos expressing membrane-targeted GFP (*UAS-mCD8GFP*) under the control of the pan-neuronal driver *elav-GAL4.* **(i–vi)** Control embryos (*elav-GAL4::UAS-mCD8GFP*) display stereotyped axonal architecture and organized neuronal networks. **(vii–xii)** Embryos expressing Aβ (*elav-GAL4::UAS-mCD8GFP > UAS-A*β) and **(xiii–xviii)** embryos expressing Tau (*elav-GAL4::UAS-mCD8::GFP > UAS-Tau*) exhibit altered axonal patterning and disrupted neuronal organization. **(B)** Representative confocal images of stage 15–16 embryos expressing *UAS-mCD8GFP* under the control of the neuroblast-specific driver *insc-GAL4*. **(i–vi)** Control embryos show well-organized axonal trajectories and intact neuronal architecture. **(vii–xii)** Aβ expressing embryos and **(xiii–xviii)** Tau-expressing embryos display pronounced defects in axonal organization and cytoskeletal integrity. GFP (green) labels neuronal membranes and projections **(i, vii, xiii),** 22C10 (red) marks Futsch, architecture **(ii, viii, xiv)**. Merged images are shown in **(iii, ix, xv)**, and merged images with DAPI nuclear staining are shown in **(iv, x, xvi)**. Higher-magnification views **(v, xi, xvii)** and **(vi, xii, xviii)** highlighted with yellow dashed outlines. Images (A-B) are maximum intensity projections of all the sections. Scale bar 50 µm.

In control embryos, Futsch expression was also clearly visible in the dendrites of the lateral chordotonal organs, reflecting normal dendritic organization. This pattern was markedly disrupted in Aβ- and Tau-expressing embryos, where dendritic labelling appeared fragmented and disorganized, and further supporting compromised neuronal cytoskeletal integrity. Notably, axonal and dendritic defects were consistently more severe when Aβ or Tau expression was driven in neuroblasts using *inscGAL4*, compared with neuronal expression alone, *elavGAL4* suggesting that early perturbation at the progenitor level exacerbates defects in neuronal architecture.

Together, these observations demonstrate that expression of Aβ or Tau during embryogenesis disrupts axonal patterning and compromises neuronal cytoskeletal organization. These early architectural defects indicate that Alzheimer’s disease associated proteins may interfere with fundamental processes of neural circuit formation during development.

### 2.2 Aβ and Tau induce irregular Prospero-mediated neuronal differentiation and impair progenitor lineage progression

During *Drosophila* neurodevelopment, neuroblasts act as neural stem cells that divide asymmetrically to self-renew while generating ganglion mother cells (GMCs), which subsequently differentiate into neurons. This lineage progression is tightly regulated by the transcription factor Prospero, which is cytoplasmically localized in neuroblasts but translocates to the nucleus in GMCs, where it represses stem cell gene expression and promotes neuronal differentiation (Li et al., 2014). Proper spatial regulation of Prospero is therefore essential for maintaining neuroblast identity and ensuring balanced neurogenesis.

To determine whether Aβ or Tau-induced neurotoxicity alters Prospero regulation, we examined its expression and localization in stage 15–16 embryos and third-instar larval brains using neuron-specific (*elav-GAL4*) and neuroblast-specific (*insc-GAL4*) drivers. In control embryos, Prospero expression was segmentally organized and largely restricted to GMCs and newly differentiating neurons within the ventral nerve cord, consistent with normal lineage specification [(Fig. 2B (i–iii); Fig. 2D (i–iii)]. In contrast, embryos expressing Aβ or Tau exhibited a marked increase in the number of Prospero-positive cells throughout the ventral nerve cord [(Fig. 2B (iv–ix); Fig. 2D (iv–ix)]. This elevation was accompanied by ectopic and enhanced Prospero expression beyond its normal lineage-restricted pattern, suggesting aberrant activation of differentiation programs. These defects were observed following both neuronal and neuroblast-specific expression, with more pronounced disruption when Aβ or Tau was expressed in neuroblasts.

**Figure 2:**
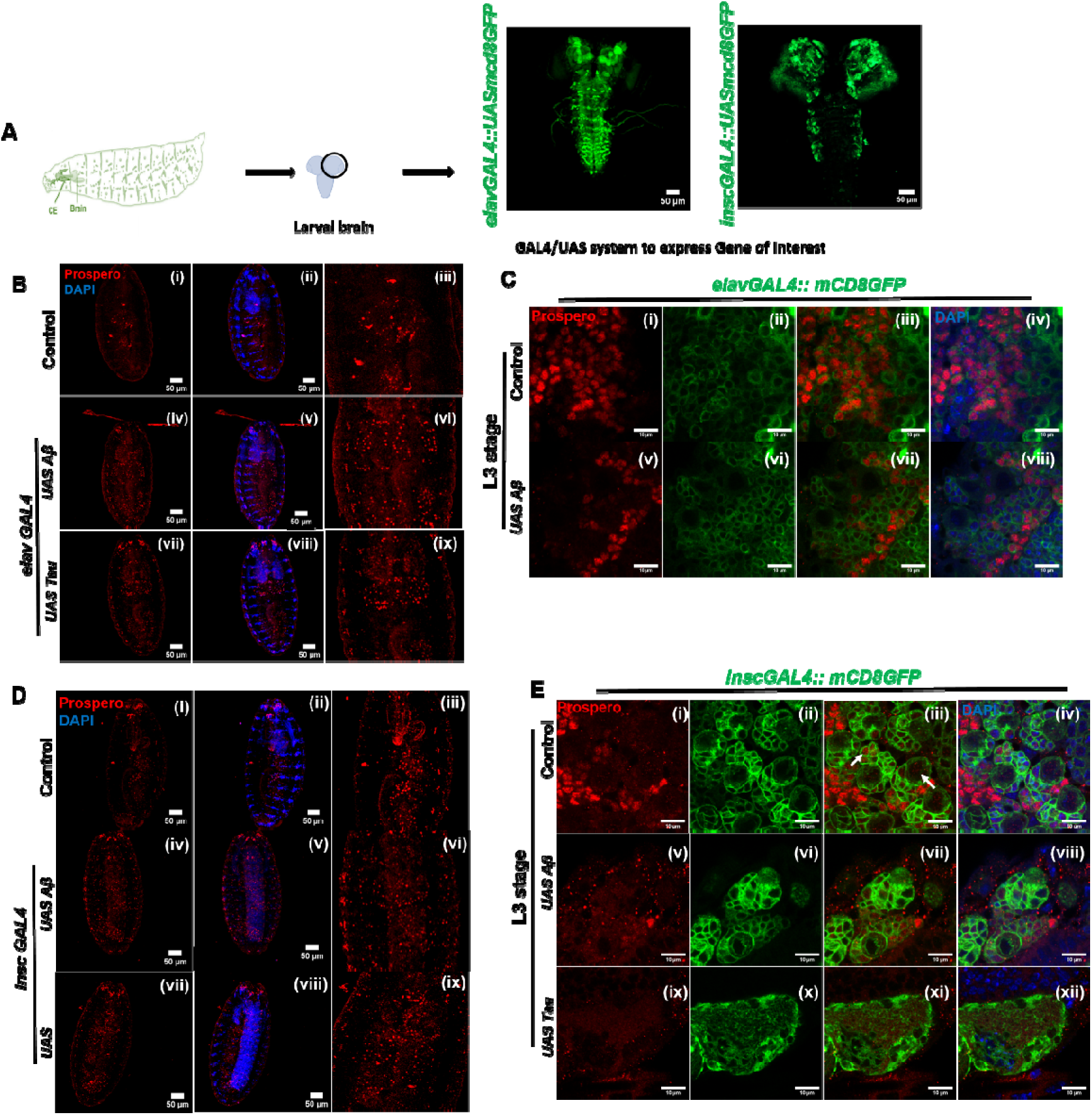
Amyloid-β and Tau induce aberrant Prospero expression in neurons and neuroblasts during embryonic and larval development. **(A)** Schematic representation of third instar (L3) larval brains using the GAL4/UAS system to drive expression of the gene of interest and visualization of larval brains expressing *UAS-mCD8GFP* under *elav-GAL4* and *insc-GAL4* drivers. **(B)** Representative confocal images of stage 15–16 embryos expressing *UAS-mCD8GFP* under the control of *elav-GAL4*, immunostained for Prospero (red) and counterstained with DAPI (blue). **(i–iii)** Control embryos show stereotyped Prospero expression. **(iv–vi)** Embryos expressing Aβ (*elavGAL4 > UAS-A*β) and **(vii–ix)** Tau (*elav-GAL4 > UAS-Tau*) display altered Prospero distribution and an increase number of Prospero-positive cells. **(C)** Confocal images of third-instar larval (L3) brains expressing *UAS-mCD8GFP* under elav-GAL4, stained for Prospero. **(i–iv)** Control larval brains show well-defined Prospero-positive lineages. **(v–viii)** Aβ-expressing larval brains exhibit disrupted Prospero patterning and reduced lineage organization. **(D)** Representative images of stage 15–16 embryos expressing *UAS-mCD8GFP* under the control of *insc-GAL4*, stained for Prospero and DAPI. **(i–iii)** Control embryos exhibit robust and spatially restricted Prospero expression in neuroblast lineages. **(iv–vi)** Aβ expressing and Tau **(vii–ix)** expressing embryos show mislocalization accompanied by an altered number of Prospero-positive cells. **(E)** Confocal images of L3 larval brains expressing *UAS-mCD8GFP* under *insc-GAL4*, immunostained for Prospero and counterstained with DAPI. **(i–iv)** Control brains display organized Prospero-positive neuroblast lineages. **(v–viii)** Aβ and Tau **(ix–xii)** expression leads to disrupted Prospero localization and altered lineage architecture. Across all panels, GFP (green) marks neuronal membranes and progenitor-derived lineages, Prospero (red), and nuclei are labeled with DAPI (blue). Images (C and E) are single optical sections while images (B and D) are maximum intensity projections of all the sections. Scale bars: 10 and 50 µm.

To assess whether Prospero misregulation persisted beyond embryogenesis, we analyzed third-instar larval brains. In control larval brains, Prospero expression was largely confined to GMCs, showing overlap with neuronal populations marked by *elavGAL4> mCD8GFP* or with neuroblast populations marked by *inscGAL4* [(Fig. 2C (i–iv); Fig. 2E (i–iv)]. In contrast, Aβ-expressing larval brains displayed aberrant Prospero localization, with decreased co-localization in neuronal populations and a subsequent reduction in Prospero-positive GMCs [(Fig. 2C (v–viii)]. Similarly, expression of Aβ [(Fig. 2E (v–viii)] and Tau [(Fig. 2E (ix–xii] in neuroblasts resulted in a reduced number of Prospero-positive GMCs and a spatially restricted neuroblast population, particularly within the central brain region (Fig. S1A, B).

Notably, Tau expression driven by *elavGAL4* resulted in severe developmental lethality, while less than 15% survival was observed when Tau was expressed using *insc-GAL4*. Owing to this lethality, Prospero analysis in Tau-expressing larval brains was limited. The pronounced toxicity is consistent with the essential role of Tau as a microtubule-associated protein required for cytoskeletal organization during mitosis, and suggests that Tau overexpression severely compromises neural lineage viability.

Collectively, these findings demonstrate that Aβ and Tau disrupt Prospero-dependent lineage regulation by inducing aberrant differentiation and impairing neuroblast lineage progression. Misregulation of Prospero at both embryonic and larval stages likely contributes to depletion of progenitor pools and defective neurogenesis during development.

### 2.3 Embryonic glial patterning is preserved, with subtle alterations in larval brains expressing Aβ and Tau

Glial cells play essential roles in nervous system development by supporting neuronal differentiation, maintaining tissue architecture, and ensuring proper neuronal–glial interactions. In the *Drosophila* embryonic nervous system, glial identity is established shortly after stage 11, following transient expression of the transcription factor *glial cells missing* (*gcm*). From stage 14 onwards, most ventral nerve cord (VNC) glia expresses the pan-glial marker *Reversed Polarity* (*repo*), allowing reliable assessment of glial patterning during late embryogenesis.

To determine whether Aβ and Tau expression affects glial development, we examined Repo expression in stage 15–16 embryos and third-instar larval brains following neuronal (*elav-GAL4*) and neuroblast-specific (*insc-GAL4*) expression of these proteins. In control embryos, Repo-positive glial cells displayed a stereotyped, bilaterally symmetric distribution along the ventral nerve cord, consistent with normal embryonic glial organization [(Fig. 3A (i–iii); Fig. 3C (i–iii)]. Surprisingly, embryos expressing Aβ and Tau exhibited largely comparable glial patterns, with no overt disruption in overall glial number or spatial distribution relative to controls [(Fig. 3A (iv–ix); Fig. 3C (iv–ix)]. Repo-positive cells maintained their characteristic positioning along the VNC, indicating that early glial specification and gross embryonic glial patterning are largely preserved despite neuronal and progenitor defects induced by Aβ and Tau.

**Figure 3:**
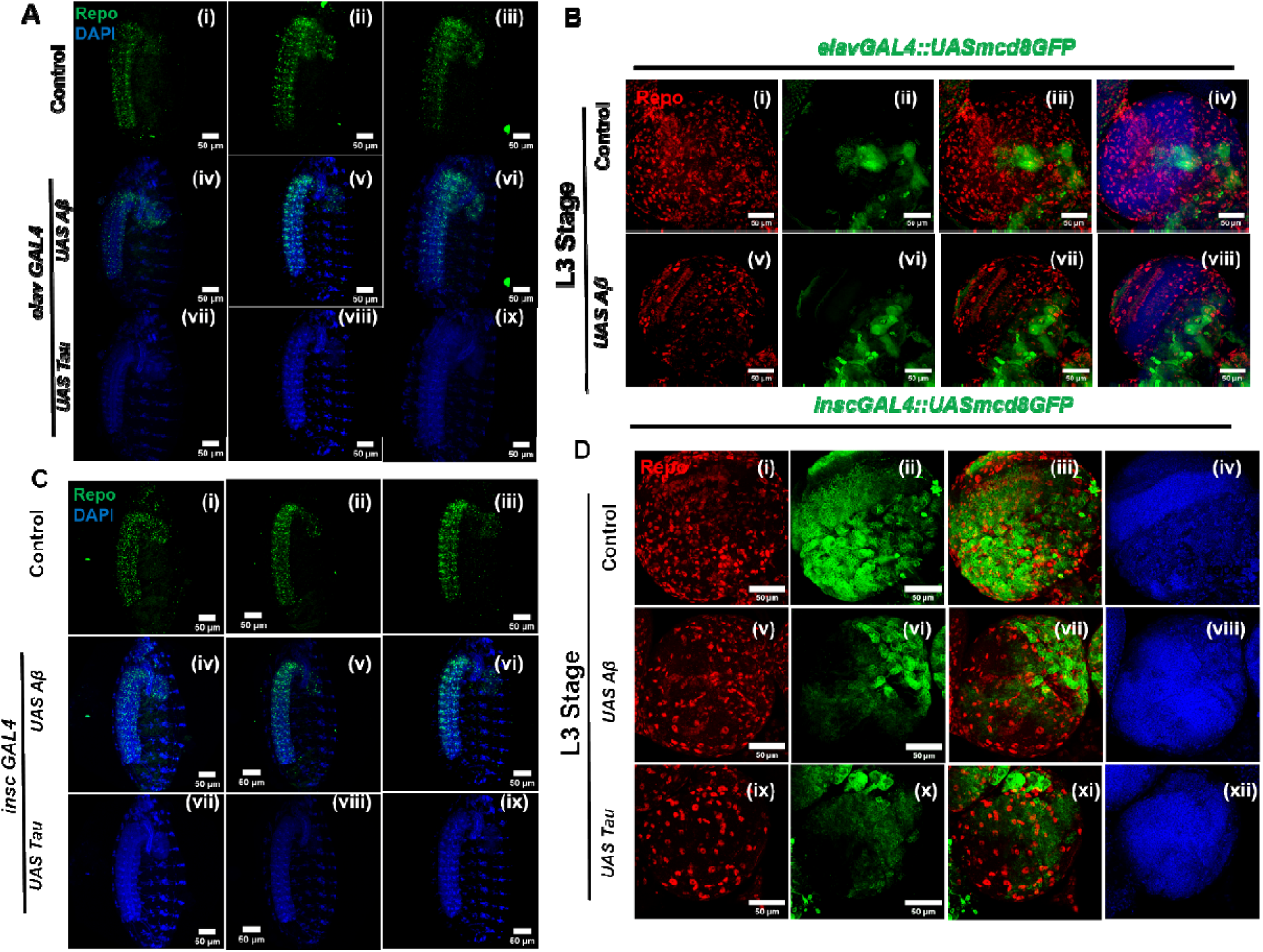
Glial identity is preserved with subtle alterations in spatial organization in Aβ- and Tau-expressing nervous systems. **(A)** Confocal images of stage 15–16 embryos stained for the pan-glial marker Repo (green) and DAPI (blue). **(i–iii)** In control embryos, Repo-positive glial cells exhibit a continuous and stereotyped distribution along the ventral nerve cord. Embryos expressing Aβ **(iv–vi)** and Tau **(vii–ix)** show a broadly comparable glial pattern, with no gross disruption of embryonic glial organization. **(B)** Confocal images of third-instar (L3) larval brains expressing *UAS-mCD8GFP* under *elav-GAL4* (green), stained for Repo (red) and DAPI (blue). **(i–iv)** Control larval brains display evenly distributed Repo-positive glial cells with well-defined associations with neuronal territories. In Aβ-expressing larval brains **(v–viii)**, glial identity and overall abundance are maintained; however, Repo-positive cells exhibit subtle spatial alterations, including reduced uniformity of spacing and localized clustering. **(C)** Representative confocal images of stage 15–16 embryos stained for Repo (green) and DAPI (blue) in control, *insc-GAL4 > UAS-A*β, and *insc-GAL4 > UAS-Tau* backgrounds. Embryonic glial distribution remains largely comparable across genotypes, indicating preservation of early glial patterning upon neuroblast-specific expression of Aβ or Tau. **(D)** Confocal images of L3 larval brains expressing *UAS-mCD8GFP* under *insc-GAL4* (green) and stained for Repo (red) and DAPI (blue). While Repo-positive glial cells remain abundant and broadly distributed in Aβ **(v–viii)** and Tau-expressing **(ix–xii)** brains, mild disorganization is evident, characterized by less uniform glial spacing and less sharply defined glial–neuronal boundaries compared with controls **(i–iv).** Images (A-D) are maximum intensity projections of all the sections. Scale bars: 50 µm.

We next examined third-instar larval brains to assess whether glial organization is affected at later developmental stages. In control larval brains, Repo-positive glial cells were abundant and uniformly distributed, forming well-defined associations with neuronal territories in both neuronal and neuroblast-specific driver backgrounds [(Fig. 3B (i–iv); Fig. 3D (i–iv)]. In contrast, larval brains expressing Aβ and Tau displayed subtle but reproducible alterations in glial cells number (Fig. S3 A) and organization. These included reduced uniformity in glial spacing, localized clustering, and less sharply defined glial neuronal boundaries, particularly in genotypes expressing Aβ and Tau under the neuroblast specific *insc-GAL4* [(Fig. 3B (v–viii); Fig. 3D (v–xii)].

Notably, Tau expression driven by *elav-GAL4* resulted in complete larval lethality, precluding analysis of glial organization in this condition. Nevertheless, the observed changes in larval glial organization in surviving genotypes expressing Aβ also showed reduced uniformity in glial spacing, localized clustering and number (Fig. S3 B), suggesting that glial alterations are likely secondary to disrupted neural architecture rather than arising from primary defects in glial development.

Together, these results indicate that Aβ and Tau do not grossly perturb early embryonic glial patterning but lead to subtle alterations in glial spatial organization during larval development, as a consequence of impaired neurogenesis and neuronal architecture.

### 2.4 Aβ and Tau expression perturb neuroblast cell-cycle dynamics in the larval brain

Proper regulation of the cell cycle in neural stem cells is essential for maintaining neuroblast self-renewal and ensuring sustained neuronal production during development. To investigate whether Aβ and Tau affects neuroblast proliferation, we analyzed cell-cycle dynamics using the **F**luorescent **U**biquitination-based **C**ell **C**ycle **I**ndicator (FUCCI) system (Zielke et al., 2014). Because *insc-GAL4* is selectively expressed in dividing neural progenitors, FUCCI reporters driven by this promoter allow precise visualization of cell-cycle phases in neuroblasts, whereas the neuron-specific driver *elav-GAL4* labels post-mitotic neurons that have exited the cell cycle. Therefore, *insc-GAL4* driven FUCCI analysis was used to specifically assess the impact of Aβ and Tau on neuroblast cell-cycle progression.

In control third-instar larval brains, FUCCI reporters revealed a heterogeneous population of actively cycling neuroblasts distributed across G1 (green), S (red), and G2/M (yellow/overlapping) phases, indicative of robust and coordinated proliferative activity [(Fig. 4B (i–iv)]. Neuroblasts were evenly distributed throughout the brain lobes, reflecting normal stem cell maintenance and lineage progression. In contrast, neuroblasts expressing Aβ exhibited a marked reduction in both the number and overall intensity of FUCCI-positive cells [(Fig. 4B (v–viii)]. This reduction was accompanied by a pronounced depletion of S-phase and G2/M populations, with a relative enrichment of G1-phase cells. These observations suggest impaired cell-cycle progression and a tendency toward delayed proliferation or premature cell-cycle exit in Aβ-expressing neuroblasts.

**Figure 4:**
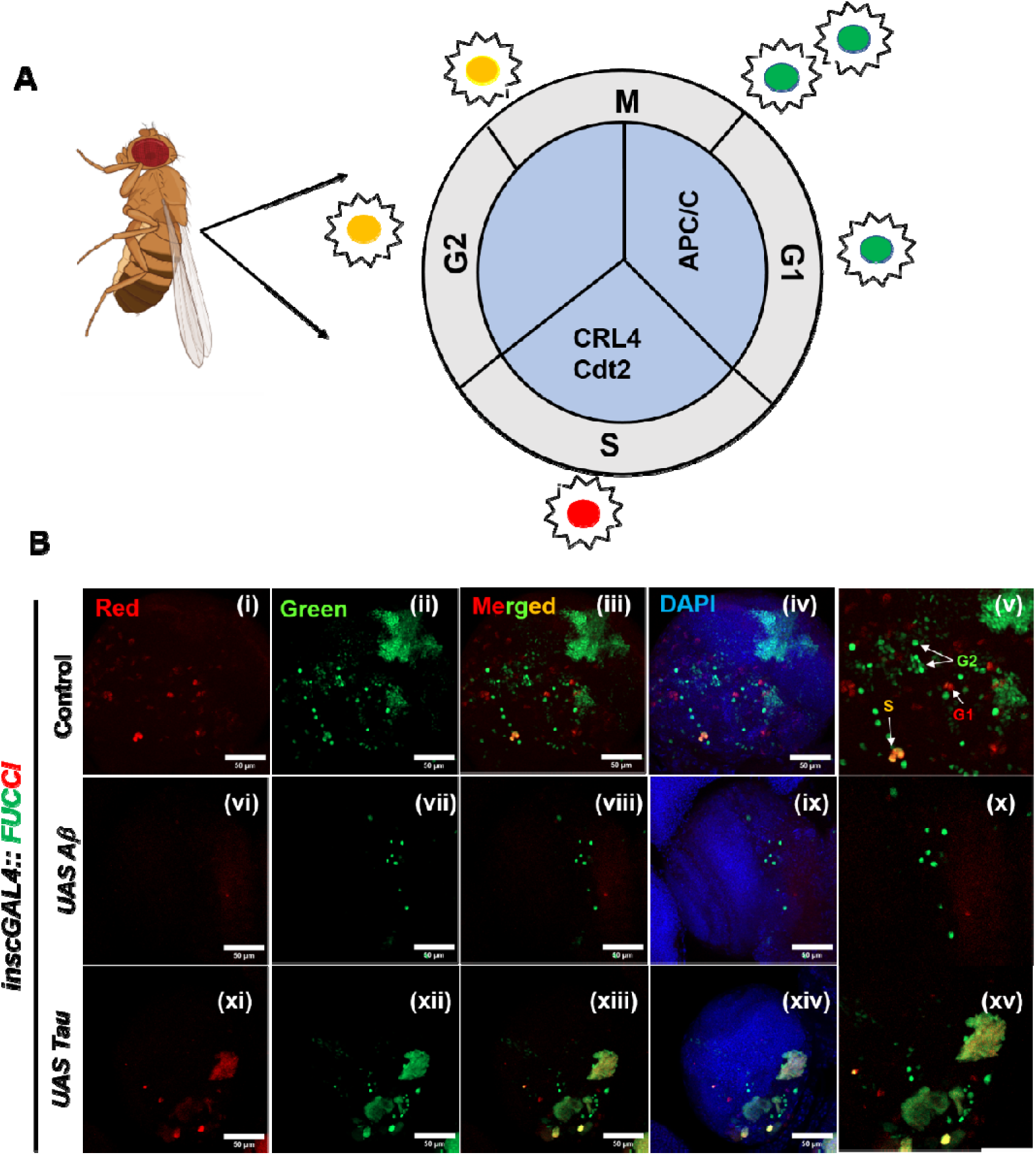
Amyloid-β and Tau disrupt neuroblast cell-cycle dynamics in the larval brain. **(A)** Schematic representation of the Fly-FUCCI (Fluorescent Ubiquitination-based Cell Cycle Indicator) system used to monitor cell-cycle progression in *Drosophila*. Cells in G1 phase are labeled in green, S-phase cells are labeled in red, and cells in G2/M phase appear yellow due to overlapping signals. Cell-cycle–dependent degradation of FUCCI reporters is regulated by APC/C and CRL4-Cdt2 E3 ubiquitin ligase complexes. **(B)** Representative confocal images of third-instar (L3) larval brains expressing the FUCCI reporters under the control of *insc-GAL4*. **(i–v)** Control brains display a balanced distribution of FUCCI-labeled neuroblasts across different cell-cycle phases, indicating normal cell-cycle progression. **(vi–x)** Neuroblast-specific expression of Aβ (*inscGAL4 > UAS-A*β) results in an altered distribution of FUCCI signals, with reduced representation of cycling cells and accumulation in specific cell-cycle phases. **(xi–xv)** Tau expression (*inscGAL4 > UAS-Tau*) similarly perturbs neuroblast cell-cycle dynamics, leading to aberrant FUCCI patterns compared with controls. Red and green channels indicate FUCCI reporters, merged images are shown alongside DAPI (blue) nuclear staining. Panel **(v)** shows magnified images of individual cell-cycle phases (G1, S, G2/M) based on FUCCI signal combinations. Images (B) are maximum intensity projections of all the sections. Scale bars: 50 µm.

Tau-expressing neuroblasts displayed a distinct but equally aberrant FUCCI pattern [(Fig. 4B (ix–xii)]. Rather than a uniform distribution of cycling cells, FUCCI-positive neuroblasts appeared clustered and exhibited heterogeneous cell-cycle signatures, suggesting disrupted coordination of cell-cycle transitions. This irregular pattern is indicative of altered or asynchronous progression through the cell cycle in response to Tau expression.

Together, these results demonstrate that both Aβ and Tau interfere with normal neuroblast cell-cycle dynamics, albeit through distinct patterns of disruption. Aβ primarily reduces proliferative capacity, whereas Tau induces disorganized and aberrant cell-cycle behaviour, likely contributing to neuroblast depletion and impaired neurogenesis during larval brain development.

### 2.5 Aβ and Tau elevate apoptotic signaling in neuronal and neuroblast lineages

During neural development, tightly regulated apoptosis plays an essential role in shaping the nervous system; however, excessive or ectopic activation of apoptotic pathways can compromise tissue integrity and lineage maintenance. Based on the pronounced defects observed in neurogenesis, cell-cycle dynamics, and neuronal architecture in Aβ and Tau-expressing genotypes, we next examined whether these alterations were associated with any change in the apoptotic cell death.

To assess apoptosis, we analyzed the expression of cleaved Caspase-3, a widely used marker of apoptotic activation, in third-instar larval brains. In control larval brains, cleaved Caspase-3 immunoreactivity was low and restricted, consistent with normal levels of developmental apoptosis [(Fig. 5A-B (i–iii)]. In contrast, expression of Aβ resulted in a pronounced increase in Caspase-3 signal intensity under both neuroblast-specific (*insc-GAL4*) and neuronal (*elav-GAL4*) drivers [(Fig. 5A–B (iv–vi)], indicating elevated apoptotic stress in neural tissues.

**Figure 5:**
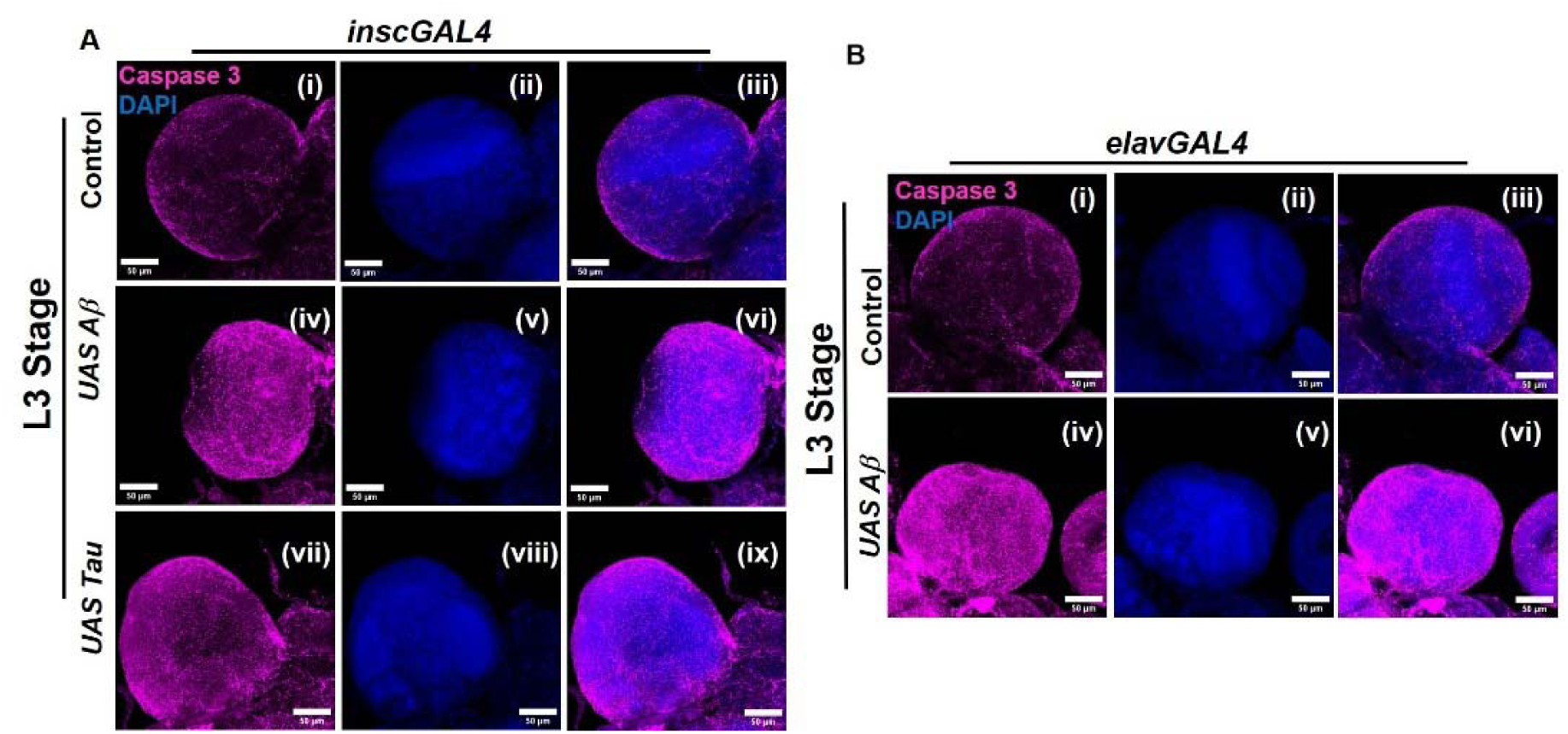
Amyloid-β and Tau elevate apoptotic signaling in larval neural tissues. **(A)** Confocal images of third-instar (L3) larval brains immunostained for cleaved Caspase-3 (magneta) and counterstained with DAPI (blue) under the control of *insc-GAL4*. **(i–iii)** Control brains show low and restricted Caspase-3 signal. **(iv–vi)** Neuroblast-specific expression of Aβ (*insc-GAL4 > UAS-A*β) results in elevated Caspase-3 immunoreactivity. (vii–ix) Tau expression *(inscGAL4 > UAS-Tau*) similarly leads to increased apoptotic signaling compared with controls. **(B)** Confocal images of L3 larval brains expressing transgenes under the pan-neuronal driver *elav-GAL4*, stained for cleaved Caspase-3 (magenta) and DAPI (blue). **(i–iii)** Control brains exhibit minimal Caspase-3 signal. **(iv–vi)** Neuronal expression of Aβ (*elavGAL4 > UAS-A*β) induces increased Caspase-3 immunoreactivity, indicating elevated apoptotic stress. Images (A-B) are maximum intensity projections of all the sections. Scale bars: 50 µm.

Similarly, Tau expression driven by *insc-GAL4* led to a marked increase in cleaved Caspase-3 levels in larval brains [(Fig. 5A (vii–ix)], suggesting enhanced apoptosis within the neuroblast lineage. Quantitative analysis confirmed a significant increase in Caspase-3 fluorescence intensity in Aβ- and Tau-expressing brains compared with controls (Fig. S4 A, B). Owing to complete larval lethality associated with Tau expression under *elav-GAL4*, apoptotic analysis could not be performed in this condition.

Notably, increased apoptotic signaling was observed in the context of reduced neuroblast numbers, impaired cell-cycle progression, and disrupted lineage specification, suggesting that apoptosis contributes to the depletion of neural progenitors and neuronal populations in Aβ-and Tau-expressing brains. These findings indicate that Alzheimer’s disease–associated proteins elicit elevated apoptotic responses during larval brain development, which may exacerbate defects in neurogenesis and brain organization.

## 3. DISCUSSION

Alzheimer’s disease (AD) is a progressive neurodegenerative disease that worsens with aging; however, increasing evidence suggests that disease-associated proteins may exert pathogenic effects long before the onset of overt neurodegeneration. The majority of studies focus on progression of disease after the onset of disease (Kamatham et al., 2024; Tahami et al., 2022) Although disease progression has been widely studied, the molecular and cellular events triggering disease onset remain largely unclear. In this study, we demonstrate that expression of Aβ and Tau during development disrupts the process of neurogenesis in *Drosophila*, including axonal architecture, neural progenitor lineage progression, cell-cycle regulation, and neuronal survival. Our finding provides empirical evidence that AD-associated proteins induced neurotoxicity viz. Aβ and Tau, compromise neurodevelopmental processes at embryonic and larval stages, supporting the emerging concept that neurodevelopmental perturbations may contribute to later neurodegenerative vulnerability.

During the embryonic development in Aβ and Tau expressing embryos in both progenitor cells viz. neuroblast and differentiated neuron, the most pronounced phenotypic observed was the disruption of axonal patterning and neuronal cytoskeletal organization. Proper axon fasciculation and commissural formation are fundamental for establishing functional neural circuits, and these processes rely heavily on intact microtubule dynamics and cytoskeletal stability (Prokop 2020; Prokop et al., 2013). The observed axonal defasciculation, discontinuities, and misrouting suggest that Aβ and Tau interfere with cytoskeletal organization during a critical window of neural circuit assembly as shown by 22C10 staining. Given the established role of Tau as a microtubule-associated protein (Tiwari and Tapadia, 2025; Vourkou et al., 2023; Melanie et al., 2022;), our findings indicate that early perturbation of cytoskeletal integrity and its impairment may represent a foundational event linking developmental defects to later neurodegeneration.

Beyond cyto-architectural defects, our data reveal profound alterations in neural stem cell lineage regulation. Prospero is a key determinant of asymmetric neuroblast division, acting as a molecular switch that promotes differentiation while restricting stem cell identity (Homem & Knoblich, 2012). The ectopic and elevated Prospero expression observed in Aβ- and Tau-expressing embryos in both neuroblast/ lineage specific and differentiated neuron. This suggests premature activation of differentiation programs, potentially at the expense of neuroblast self-renewal. However, larval brains displayed a reduction in Prospero-positive ganglion mother cells and a spatially restricted neuroblast population, implying the impaired lineage progression. These observations align with previous studies reporting that dysregulation of fate determinants can lead to stem cell exhaustion and reduced neuronal output, and extend this concept to models of Alzheimer’s disease–associated protein toxicity (Meyer et al., 2019).

As the similar defects in cytoskeletal integrity and prospero positive cells distribution, were observed when Aβ and Tau are expressed in differentiated neurons and neuroblast, we used the Fly-FUCCI system to understand the cell cycle dynamics during third instar stage only. Impaired neuroblast cell-cycle dynamics further supports the notion that Aβ and Tau compromise neural progenitor maintenance. The genetic analysis using the Fly-FUCCI system revealed that Aβ expression leads to a depletion of proliferative neuroblasts, accompanied by enrichment of cells in the G1 phase, indicative of delayed cell-cycle progression or premature cell-cycle exit. In contrast, Tau expression resulted in a heterogeneous and disorganized cell-cycle pattern, suggesting aberrant regulation of cell-cycle transitions. Although Aβ and Tau appear to perturb neuroblast proliferation through distinct mechanisms, both ultimately converge on reduced proliferative capacity and impaired neurogenesis. Such defects in stem cell dynamics during development could have lasting consequences for brain structure and function (Homem & Knoblich, 2012).

Interestingly, glial organization remained largely undisturbed during embryogenesis, with only subtle spatial alterations observed at larval stages, in both neuron and neuroblast specific expression of Aβ and Tau. This relative resistance of glial patterning to early Aβ and Tau toxicity suggests that neurons and neural progenitors are more vulnerable during initial developmental stages (Rahman et al., 2022; Stevenson et al., 2020). The gross glial spatial organization disruption observed later likely reflects secondary consequences of disrupted neuronal architecture and reduced neuronal populations, rather than primary defects in glial specification. These findings highlight the selective sensitivity of neuronal lineages to AD-associated proteins during development.

Increased apoptotic signaling in larval brains expressing Aβ or Tau further contributes to the observed depletion of neural cells. Despite elevated cleaved Caspase-3 expression in disease background, overall brain architecture remains intact, implying enhanced apoptotic signaling rather than widespread cell loss. Notably, Tau expression resulted in severe developmental lethality, particularly when driven in neurons, highlighting the potent toxicity of Tau during development. Together, these results suggest that excessive apoptosis acts in concert with impaired proliferation and premature differentiation to reduce neural cell numbers and disrupt brain organization.

Collectively, our study supports a model in which amyloid-β and Tau interfere with fundamental neurodevelopmental processes, including cytoskeletal organization, asymmetric stem cell division, cell-cycle regulation, and neuronal survival. Rather than acting solely as later stage neurotoxic agents, these proteins appear capable of compromising nervous system development, potentially priming neural circuits for increased susceptibility to degeneration later in life. While *Drosophila* lacks the complex cognitive phenotypes associated with human Alzheimer’s disease, the conservation of core developmental and cellular mechanisms highlights the relevance of these findings.

Future studies will be required to determine whether early developmental defects induced by Aβ and Tau lead to long-term functional consequences in adult neural circuits, and to dissect the molecular pathways linking cytoskeletal disruption to stem cell misregulation. Understanding how neurodevelopmental perturbations intersect with neurodegenerative processes may offer new perspectives on disease etiology and highlight early intervention windows for Alzheimer’s disease.

## 4. MATERIALS AND METHODS

### 4.1 *Drosophila* stocks and husbandry

*Drosophila melanogaster* stocks were maintained on a standard laboratory diet containing 10% sucrose, 10% yeast, 2% agar, 10% autolyzed yeast, supplemented with 3% nipagin and 0.3% propionic acid. Flies were reared at 25 ± 1°C under controlled conditions with 50–70% relative humidity and a 12 h light/12 h dark cycle. The following fly stocks were used in the experiment: *inscutable-GAL4 (insc),* also known as *1407-GAL4* (a generous gift from Dr. J. A. Knoblich, Vienna, Austria) recombined with *UAS-mCD8GFP* (BL-5137), *UAS-FUCCI* (BL-55122), *UAS-tau–lacZ* (BL-5829), *elavGAL4::UASmCD8GFP* (BL-5146), *elav-GAL4* (BL-458) and *UAS-A*β (BL-33774). The following genotypes combinations are generated in the lab for the study: *insc-GAL4::UAS mCD8GFP*, *insc-GAL4; UAS-FUCCI*.

For all the experiments, brains from wandering third-instar larvae were dissected and used, while stage 16 embryos were processed for whole-mount immunostaining. Genetic crosses were set up using 3–4-day-old virgin females and age-matched males collected, shortly after eclosion.

### 4.2 *Drosophila* brain dissection and immunostaining

Brains were dissected from healthy wandering third-instar larvae (118–120 h after hatching) in 1× phosphate-buffered saline (PBS). Dissected tissues were fixed in 4% paraformaldehyde for 30 min at room temperature, followed by washing in 0.1% PBST (1× PBS containing 0.1% Triton X-100). Samples were then incubated in a blocking solution containing 0.1% Triton X-100, 0.1% bovine serum albumin (BSA), 10% fetal calf serum (FCS), 0.1% sodium deoxycholate, and 0.02% thiomersal for 2 h at room temperature. Following blocking, tissues were incubated with primary antibodies overnight at 4°C. Samples were subsequently washed three times with 0.1% PBST for 15 min each. Appropriate fluorophore-conjugated secondary antibodies were applied for 2 h at room temperature, together with DAPI (1 µg/ml; Thermo Fisher Scientific, Cat# D1306) for nuclear counterstaining. After additional washes with 0.1% PBST, tissues were mounted in DABCO antifade mounting medium (Sigma-Aldrich, Cat# D27802) for imaging. The following stains and antibodies were used:

**Stains** : DAPI (1 mg/ml, Thermo Fisher Scientific, Cat# D1306)

**Primary Antibodies**: Anti-22C10 (DSHB, 1:50, Cat # 528403), Anti-Prospero (DSHB,1:50, Cat # 528440), Anti-Repo (DSHB, 1:10, Cat # 528448), Anti-Caspase 3 (CST, 1:100, Cat # 9662) Anti-GFP (Invitrogen,1:800, Cat # A10262). The Secondary antibodies used are as follows: donkey anti-mouse Alexa Fluor 546 (Invitrogen, 1:200) goat anti-rabbit Alexa Fluor 647 (Invitrogen, 1:200), donkey anti-chicken Alexa Fluor 488 (Invitrogen, 1:200).

### 4.3 Embryo Collection and Preparation

Adult female flies were allowed to lay eggs on agar food plates for 12–24 h at 24 ± 1°C. Collected embryos were transferred to microcentrifuge tubes, rinsed thoroughly with distilled water, and dechorionated by brief incubation in a 1:1 dilution of commercial bleach and distilled water. Following dechorionation, embryos were fixed using a 1:1 mixture of n-heptane and 4% formaldehyde prepared in 1× phosphate-buffered saline (PBS), with gentle agitation for 20 min at room temperature. After fixation, the lower aqueous phase was carefully removed, and 2 ml of 100% methanol was added to the samples, followed by vigorous shaking for ∼20 s to induce devitellinization. The upper phase containing non-devitellinized embryos and empty vitelline membranes was discarded, while the devitellinized embryos that settled at the bottom were retained and stored in methanol for subsequent immunohistochemical analyses, as previously described (Tiwari and Tapadia, 2025).

### 4.4 Larval synchronization and collection

For larval synchronization, adult flies were allowed to lay eggs on food plates for 12 h. Following egg collection, embryos were incubated at 25 °C for 12 h, after which hatched first-instar larvae were gently removed using a fine paintbrush, leaving behind unhatched embryos. The remaining embryos were further incubated for 30 min at 25 °C, and newly hatched larvae were then transferred to fresh vials containing standard laboratory food and maintained at 25 °C. Synchronized larvae were reared until the wandering third-instar stage (approximately 118–120 h after egg laying). At this stage, larval brains were dissected and processed for immunostaining as described in the Immunostaining section.

### 4.5 Microscopy and image processing

All samples were imaged using a Zeiss LSM-780 (Imager.Z2) and LSM-900 confocal microscope with Zen software (version 3.4), under a 20x and 63x objective (zoom 0.8,1.4 and 1.9). A 2.5 μm optical section interval was consistently applied unless otherwise specified in the figure legend. To ensure non-biased imaging and accurate intensity comparison, identical confocal settings (laser power, gain, and offset) were used for all experimental and control samples acquired on the same day. For experiments imaged across different days, wild-type samples were used to define baseline settings, which were then applied uniformly to all genotypes within that batch. Images showing any signs of oversaturation were excluded from further analysis. Post-acquisition image processing and quantification were performed using ImageJ software (NIH, USA; https://imagej.nih.gov/ij/). Figures were assembled using Adobe Photoshop 2021 (version 22.4.2), and schematic models were created using Microsoft Powerpoint.

### 4.6 Image quantification and statistical analysis

All quantification was done using ImageJ software (NIH, USA). The number of Repo positive cells was counted manually and analyzed separately. To determine the mean fluorescent intensity of Caspase 3 staining, the single ROI (80*80 μm square ROI) of the brain lobes of the maximum intensity projection image was utilized. Graphs were plotted using GraphPad Prism 7.0. Statistical analyses were performed using either a two-tailed unpaired Student’s *t*-test (for comparison between two groups) or one-way ANOVA followed by Tukey’s post-hoc test (for comparisons among multiple groups). Data are represented as mean ± standard error of the mean (SEM), with three biological replicates (n = 3). The significance level is indicated by an ∗ for *p* ≤ 0.05, ∗∗ for *p* ≤ 0.01, ∗∗∗ for *p* ≤ 0.001, ∗∗∗∗ for *p* ≤ 0.0001, and by ns for not significant.

## Acknowledgements

We thank Bloomington *Drosophila* Stock Center and Dr. J. A. Knoblich, Vienna, Austria for sharing their fly stocks.We acknowledge Interdisciplinary School of Life Sciences and Dr. Bama Charan Mondal, for confocal facility. Neha Tiwari was supported by a research fellowship from the University Grant Commission, New Delhi, and is highly acknowledged. Khushboo Sharma and Madhu G. Tapadia thank Institute of Eminence, Banaras Hindu University, Varanasi.

