## Supplementary Data for "Early disruption of neurogenesis and neural architecture by Amyloid-β and Tau during *Drosophila* development"

### Supplementary material:

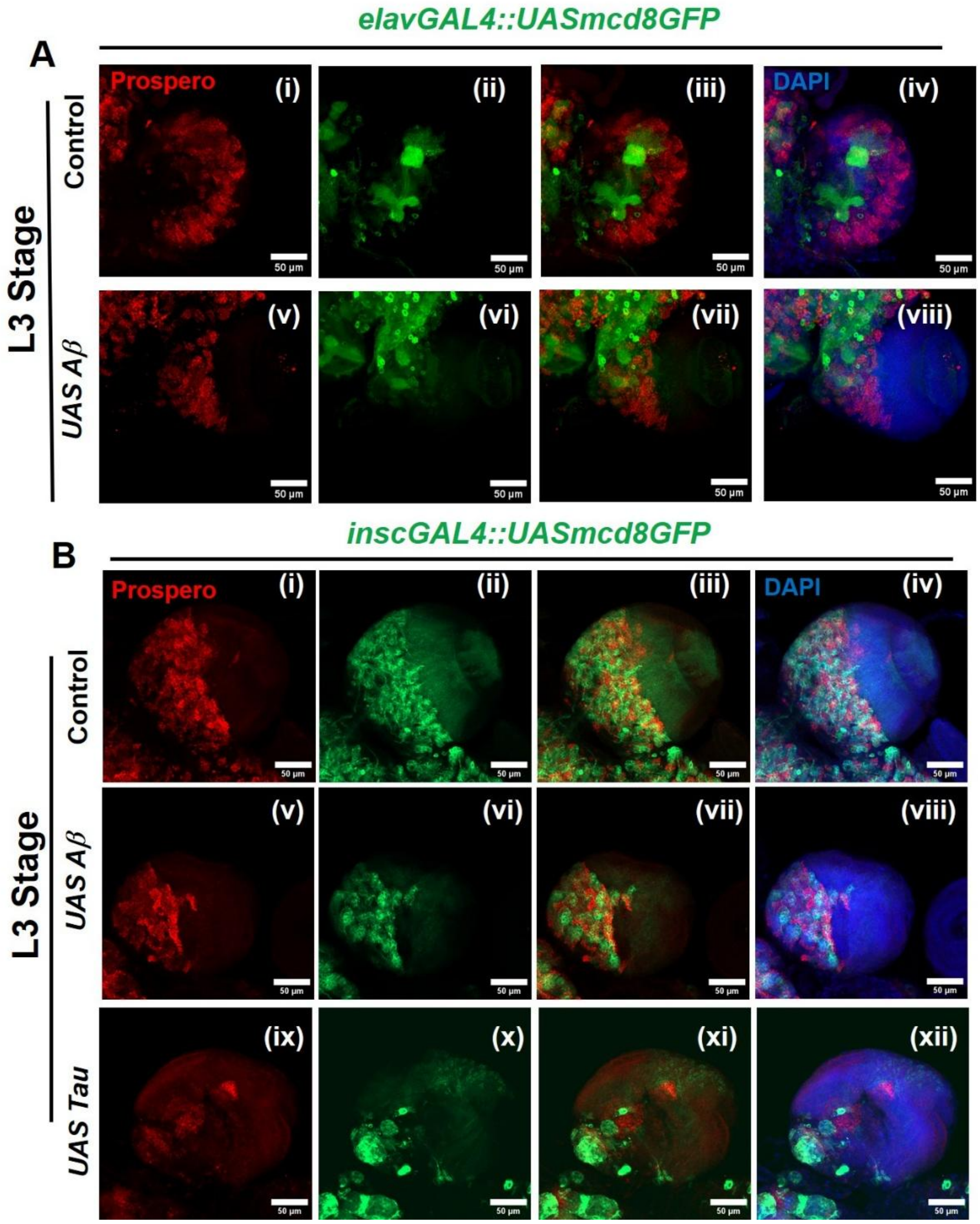


**Figure S1: Aβ and Tau expression alters Prospero distribution and neuroblast lineage organization in larval brains.**

Confocal images of third-instar (L3) larval brain lobes stained for Prospero (red), *mCD8::GFP* (green), and DAPI (blue). **(A)** Neuronal lineage labeling using *elav-GAL4 > UAS-mCD8::GFP*. In control brains **(i–iv)**, Prospero shows a characteristic pattern marking differentiating ganglion mother cells and neurons. In Aβ-expressing brains **(v–viii)**, Prospero-positive cells are still present but exhibit altered spatial distribution, with changes in clustering relative to neuronal domains.

**(B)** Neural stem cell lineage labeling using *insc-GAL4 > UAS-mCD8::GFP*. Control brains **(i–iv)** display a stereotyped arrangement of Prospero-positive cells adjacent to insc-positive neuroblast lineages. In Aβ-expressing brains **(v–viii)**, Prospero staining remains detectable but shows disrupted spatial organization and altered abundance of Prospero-positive cells. In Tau-expressing brains **(ix–xii)**, Prospero-positive cells exhibit pronounced mislocalization and changes in the number of Prospero-positive cells, indicating perturbed neuroblast lineage progression and differentiation dynamics.

Images (A-B) are maximum intensity projections of all the sections. Scale bars: 50 µm.

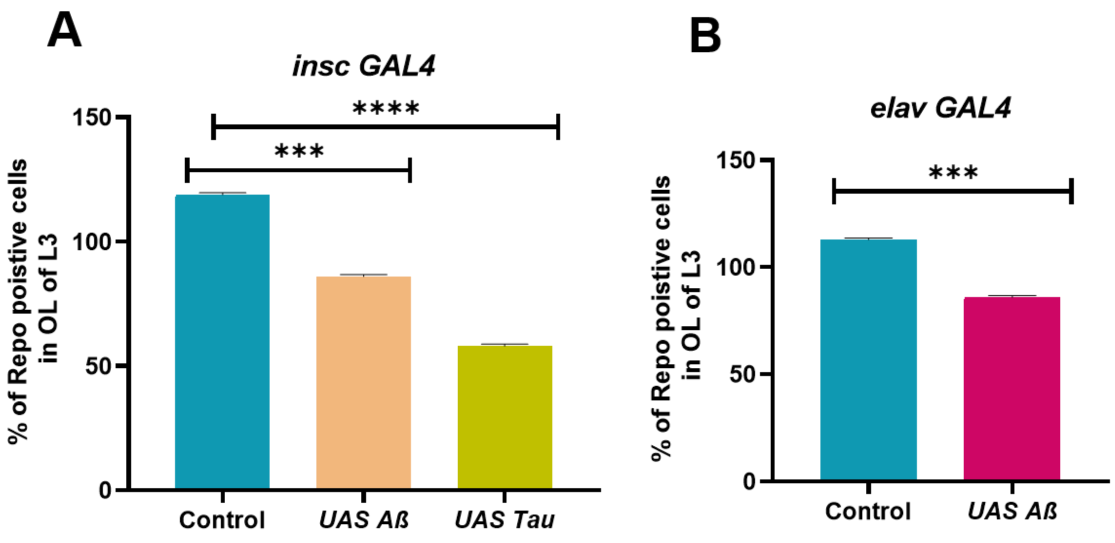


**Figure S2: Loss of Aβ and Tau reduces the number of Repo-positive glial cells in the optic lobe (OB) of L3 larval brains.**

**(A)** Quantification of Repo-positive cells in third instar larval (L3) brains driven by *insc-GAL4*. Compared to control, expression of *UAS-Aβ* and *UAS-Tau* significantly reduced the number of Repo-positive cells. **(B)** Quantification of Repo-positive cells in L3 brains driven by *elav-GAL4*, showing a significant reduction in Repo-positive cells upon *UAS-Aβ* expression compared to control.

Data represent mean ± SEM from $n \geq 10$ brains per genotype. Statistical significance was determined using one-way ANOVA (A) or unpaired two-tailed Student's *t*-test **(B)**. ***$p < 0.001$, ****$p < 0.0001$.

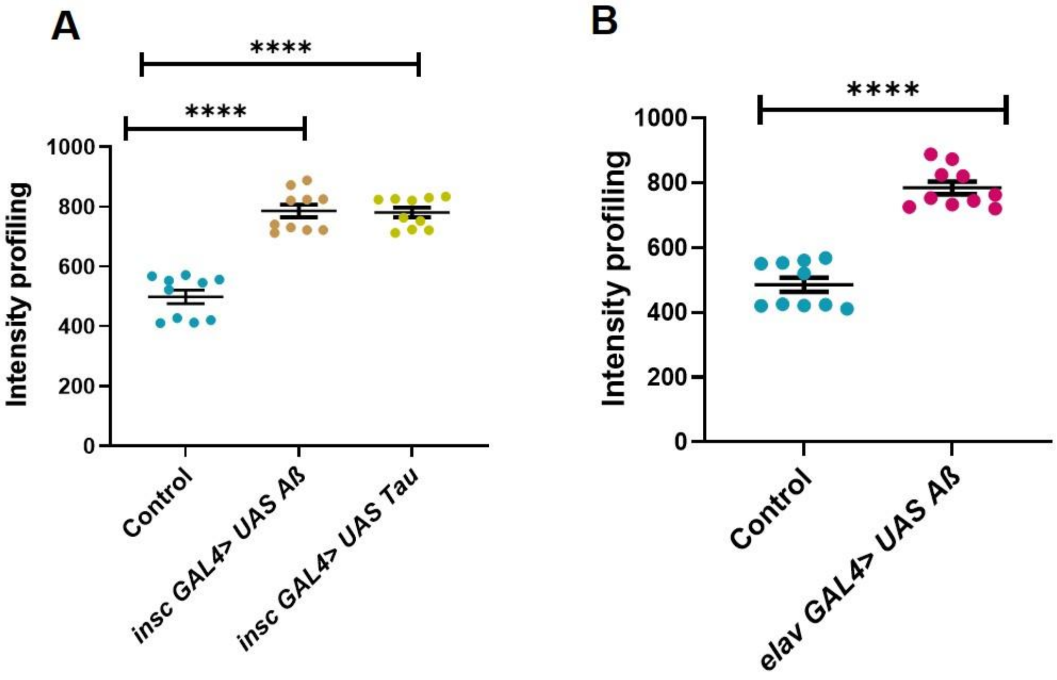


**Figure S3: Increased Caspase-3 activation upon Aβ and Tau expression.**

**(A)** Quantification of fluorescence intensity of Caspase-3 in L3 larval brains expressing *UAS-Aβ* and *UAS-Tau* under *insc-GAL4* control. Both Aβ and Tau expression significantly increased Caspase-3 levels compared to control.

**(B)** Quantification of Caspase-3 fluorescence intensity in L3 larval brains expressing *UAS-Aβ* under *elav-GAL4* control, showing a significant increase relative to control.

Each dot represents an individual brain. Horizontal bars indicate mean ± SEM. Statistical analysis was performed using one-way ANOVA (A) or unpaired Student's *t*-test (B). ****$p < 0.0001$.
